# Structural modeling of the flagellum MS ring protein FliF reveals similarities to the type III secretion system and sporulation complex

**DOI:** 10.1101/023564

**Authors:** Julien Bergeron

**Affiliations:** Department of Biochemistry and Molecular Biology, and Centre for Blood Research, University of British Columbia, Vancouver, Canada

## Abstract

The flagellum is a large proteinaceous organelle found at the surface of many bacteria, whose primary role is to allow motility through the rotation of a long extracellular filament. It is an essential virulence factor in many pathogenic species, and is also a priming component in the formation of antibiotic-resistant biofilms. The flagellum consists of the export apparatus and stator in the cytosol; the basal body, spanning the bacterial membrane(s) and periplasm; and the hook-filament, that protrudes away from the bacterial surface. Assembly of the bacterial flagellum is initiated by the formation of the basal body MS ring region, constituted of multiple copies of the protein FliF. Here, I report an analysis of the FliF sequence from various bacterial species, demonstrating that its periplasmic region is composed of a domain homologuous to that of the type III secretion system proteins PrgK, and of a second globular domain that possesses a similar fold to that of the sporulation complex component SpoIIIAG. I also describe that Chlamydia possesses an unusual FliF protein, lacking part of the PrgK homology domain and the SpoIIIAG-like domain, and fused to FliG at its C-terminus. Finally, I have combined the sequence analysis of FliF with the EM map of the MS ring, to propose the first atomic model for the FliF oligomer. These results further emphasize the similarity between the flagellum, T3SS and sporulation complex, and will facilitate further structural studies.

## INTRODUCTION

Bacteria interact with their environment using a range of surface appendages, including flagella, pili, fimbriae, and secretion systems^1^. In particular, the flagellum is responsible for motility in many bacteria^2^, but it is also frequently associated with adhesion to surfaces and/or other cells^3^. Flagella are found in many bacterial families, including most gram-positive, proteobacteria and spirochetes^4,5^. Notably, it is an essential virulence factor in many pathogenic species, such as *Salmonella, E. coli, Clostridium, Pseudomonas, Helicobacter, Vibrio, Burkholderia,* and *Campylobacter,* making the flagellum a potential target for new antibacterial therapeutics^6^.

The bacterial flagellum is constituted of four distinct regions^5,7,8^ (Figure 1A). On the cytosolic side, the export apparatus and stator are responsible for the assembly and rotation of the system. The membrane-embedded structure that traverses the cytoplasmic membrane and periplasmic space (as well as the outer membrane for gram-negative bacteria) is called the basal body. The hook is a curved filament, ~ 50 nm in length (for the prototypical *Salmonella typhymurium* flagellum), that protrudes away from the basal body. It is prolonged by the filament, a long structure (up to several μm) responsible for motility and adherence.

**Figure 1:**
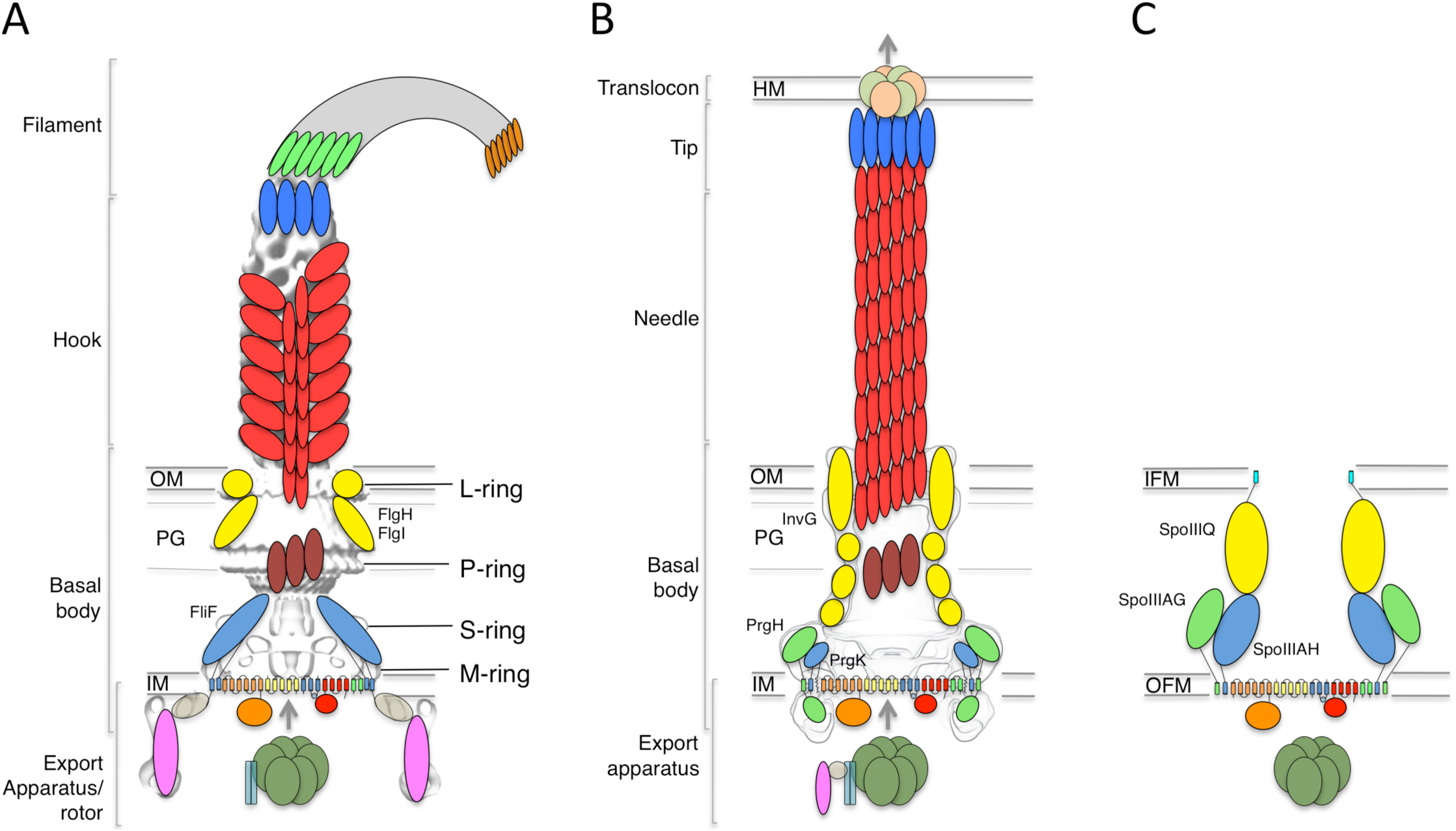
The flagellar, T3SS and sporulation complexes. Schematic representation of the bacterial flagellum (A), the T3SS (B), and the sporulation complex (C). The PrgK-like components are in blue, PrgH-like components in green, and outer-membrane components in yellow. The EM maps are shown in grey for (A) and (B). The ring structures identified in the flagellum are also indicated. IM is for inner membrane, OM is for outer membrane, IFM is for inner forespore membrane, OFM is for outer forespore membrane.

Genetic studies have revealed that the inner-membrane protein FliF is the first component of the flagellum to assemble, forming two ring-shaped structures called the MS rings^9^. This in turn recruits elements of the stator and secretion apparatus in the cytosol, leading to a secretion-competent complex, which can export the hook protein to the cytoplasm through its central pore. These observations led to the proposal that FliF is the flagellum assembly platform^10^.

FliF is an ~ 60 kDa protein, localized to the inner-membrane through the Sec pathway. Sequence analysis has indicated that it possesses two transmembrane helices, flanking a large periplasmic region^11^. At the C-terminus, a cytosolic peptide has been shown to interact with FliG^12,13^, a component of the flagellum stator. EM reconstructions of purified FliF revealed a homo 26-mer forming the MS ring oligomer^14^, although analyses of intact flagellum particles have identified 24-, 25- and 26-fold symmetry for this region of the basal body^15^.

A number of components from the cytosolic export apparatus are homologuous to that of the type III secretion system (T3SS, Figure 1B), another bacterial complex whose role is to inject so-called “effector” proteins inside the cytosol of target or symbiotic cells^16^. Indeed, phylogenetic studies have revealed that the flagellum export apparatus is likely the evolutionary ancestor of the T3SS^17^.

In particular, FliF shows significantly sequence homology to an inner-membrane component of the T3SS (24 % sequence homology with the *Salmonella* homologue PrgK, 22 % sequence identity with the EPEC homologue EscJ, for residues 52-217 of FliF)^11^. Both EscJ and PrgK form a 24-mer ring structure in the inner-membrane, similar to that of FliF, and structural characterization have revealed the molecular details of their architecture and oligomerization^18,19^. Specifically, the periplasmic region of EscJ and PrgK consists of two globular domains with a canonical “ring building motif” (RBM) fold^20^, found in several oligomeric proteins. The two RBMs are joined by a rigid linker, which was shown in PrgK to promote oligomerization^19^.

Recently, two proteins essential for the sporulation process in *Bacillus subtilis,* SpoIIIAH and SpoIIIAG, were shown to be homolgous to PrgK and FliF, ^21,22^. This led to the suggestion that these proteins are part of a complex that directs the transport of proteins and/or nutrients between the mother cell and the endospore (Figure 1C), although such complex has not been observed directly^23^.

Here, I use computational methods to demonstrate that the periplasmic region of FliF includes a PrgK homology domain, as well as a FliF-specific domain, that is similar to that of the sporulation complex component SpoIIIAG. I also report that in the Chlamydiacae family, the FliF protein differs significantly from other species, as it includes only one RBM of the PrgK homology region, and is fused to a FliG-like domain at the C-terminal cytosolic end. Finally, I combine previously determined EM maps and structural modeling to propose the first molecular model for the FliF periplasmic region, revealing that FliF is akin to a fusion of the T3SS basal body inner-membrane components PrgK and PrgH. This unexpected observation has implications in the understanding of the evolutionary relationship between the flagellum, T3SS and sporulation complex.

## RESULTS AND DISCUSSION

### Characterization of the FliF domain organization

In order to identify conserved features, I gathered FliF sequences from a number of human pathogens, spanning gram-negative, gram-positive and spirochete bacteria. Most of the FliF sequences are similar in length (~ 560 amino acids), but show limited sequence conservation (Table 1). I next used sequence analysis servers to predict secondary structure elements and other structural and functional features. Two transmembrane (TM) helices are predicted, between residues 20–45 and residues 445–470, and a secretion signal peptide targeting it for secretion and inner-membrane localization^24^ is predicted at the N-terminus. All FliF sequences show a very similar secondary sequence prediction pattern (Supplementary figure 1), and similarity to PrgK was identified for residues 50–220 in all sequences (hereafter referred to as the PrgK homology domain), but no structural homologues were found for residues 220–440 (FliF-specific domain). Within this region, residues 305–360 of the FliF-specific domain are predicted unstructured in all sequences.

**Table 1:** Sequence identity between FliF orthologues used in the multiple sequence alignment shown in Figure 2.

|  | Salmonella | E.coli | Yersinia | Bordetella | Pseudomonas | Legionella | Helicobacter | Campylobacter | Listeria | Streptococcus | Vibrio | Bacillus | Clostridium | Treponema |
| --- | --- | --- | --- | --- | --- | --- | --- | --- | --- | --- | --- | --- | --- | --- |
| Salmonella | 100.0 | 85.9 | 62.4 | 45.1 | 34.1 | 34.7 | 30.7 | 29.4 | 23.6 | 21.7 | 27.9 | 24.6 | 26.2 | 21.5 |
| E.coli |  | 100.0 | 62.6 | 46.2 | 33.7 | 34.7 | 31.4 | 29.6 | 22.0 | 22.0 | 27.7 | 25.1 | 25.0 | 22.2 |
| Yersinia |  |  | 100.0 | 49.2 | 35.5 | 34.5 | 29.6 | 28.1 | 21.2 | 21.1 | 26.6 | 24.2 | 23.9 | 19.5 |
| Bordetella |  |  |  | 100.0 | 35.9 | 35.1 | 29.6 | 26.3 | 22.9 | 21.0 | 27.5 | 23.2 | 22.0 | 21.6 |
| Pseudomonas |  |  |  |  | 100.0 | 38.1 | 28.4 | 29.0 | 20.3 | 20.3 | 31.0 | 20.0 | 22.2 | 20.3 |
| Legionella |  |  |  |  |  | 100.0 | 28.8 | 26.5 | 20.2 | 20.2 | 29.3 | 21.2 | 25.3 | 20.2 |
| Helicobacter |  |  |  |  |  |  | 100.0 | 43.2 | 23.7 | 21.9 | 23.3 | 23.8 | 28.3 | 24.2 |
| Campylobacter |  |  |  |  |  |  |  | 100.0 | 22.8 | 21.2 | 26.1 | 23.1 | 23.3 | 25.0 |
| Listeria |  |  |  |  |  |  |  |  | 100.0 | 37.4 | 19.8 | 22.3 | 24.5 | 20.5 |
| Streptococcus |  |  |  |  |  |  |  |  |  | 100.0 | 20.5 | 22.4 | 24.4 | 21.3 |
| Vibrio |  |  |  |  |  |  |  |  |  |  | 100.0 | 19.1 | 21.8 | 18.6 |
| Bacillus |  |  |  |  |  |  |  |  |  |  |  | 100.0 | 24.6 | 23.2 |
| Clostridium |  |  |  |  |  |  |  |  |  |  |  |  | 100.0 | 20.7 |
| Treponema |  |  |  |  |  |  |  |  |  |  |  |  |  | 100.0 |

I then generated a multiple alignment of all FliF sequences, and mapped the predicted secondary structure and identified domains (Figure 2). This further illustrates the conserved domain organization in all FliF orthologues. The two RBMs of the PrgK homology domain (labeled RBM1 and RBM2) are well conserved, as is the linker L1 between these. It has been shown that in PrgK this linker plays a role in ring assembly^19^, suggesting that this may also the case in FliF. In contrast, the linker region L2, separating the PrgK homology domain to the FliF-specific domain, is highly variable.

**Figure 2:**
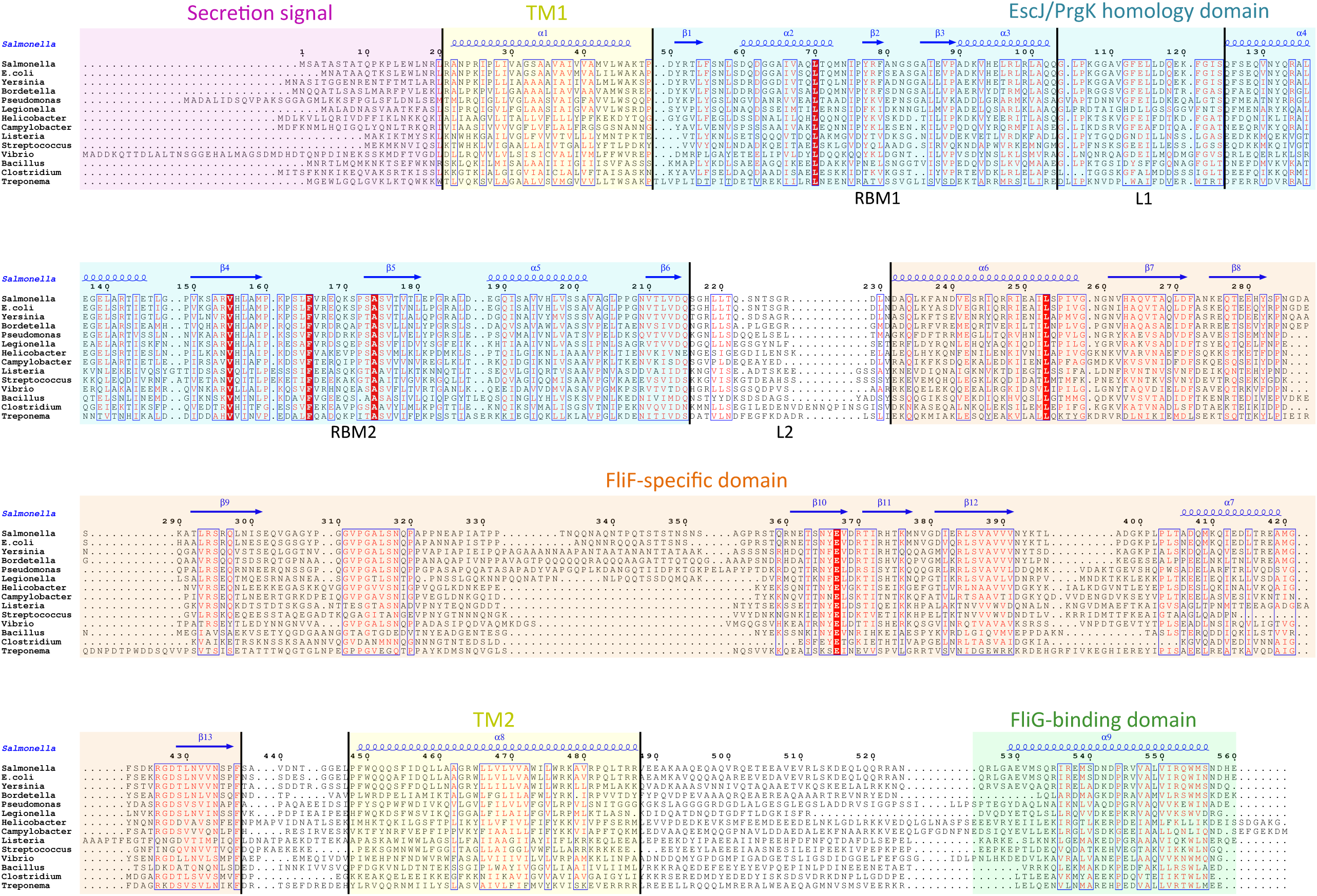
Domain organization of FliF. Multiple sequence alignment of FliF sequences from various human pathogens (*Salmonella typhimurium, Escherichia coli, Yersinia pestis, Bordetella pertussis, Pseudomonas aeruginosa, Legionella pneumophilia, Helicobacter pylori, Campylobacter jejuni, Listeria monocytogenes, Streptococcus pneumonia, Vibrio cholerae, Bacillus subtilis, Clostridium difficile, Treponema palladium*). Conserved residues are in red box, similar residues are in red characters. Identified domains are highlighted in colored boxes, with the TM helices in yellow, the FliG-binding domain in green, the signal sequence in purple, the PrgK homology domain in blue and the FliF-specific domain in orange. The predicted secondary structure elements for the *S. typhimurium* FliF are in blue at the top.

### The Chlamydiacae FliF-FliG fusion protein

While the FliF domain organization described above was found in most FliF sequences, one notable exception was identified, for the Chlamidiacae family, where the FliF sequence is notably shorter (~330 amino acids). Sequence analysis and multiple sequence alignment from all available Chlamydiacae homologues (Figure 3A) revealed that the N-terminal signal sequence and the two TM (residues 16–33 and 250–275) are present, but the periplasmic region is significantly shorter. Residues 60–145 were identified as structurally similar to PrgK, but only encompassing RBM2. No structural homologues could be identified for residues 165–235, however the predicted secondary structure matches that of the canonical RBM.

**Figure 3:**
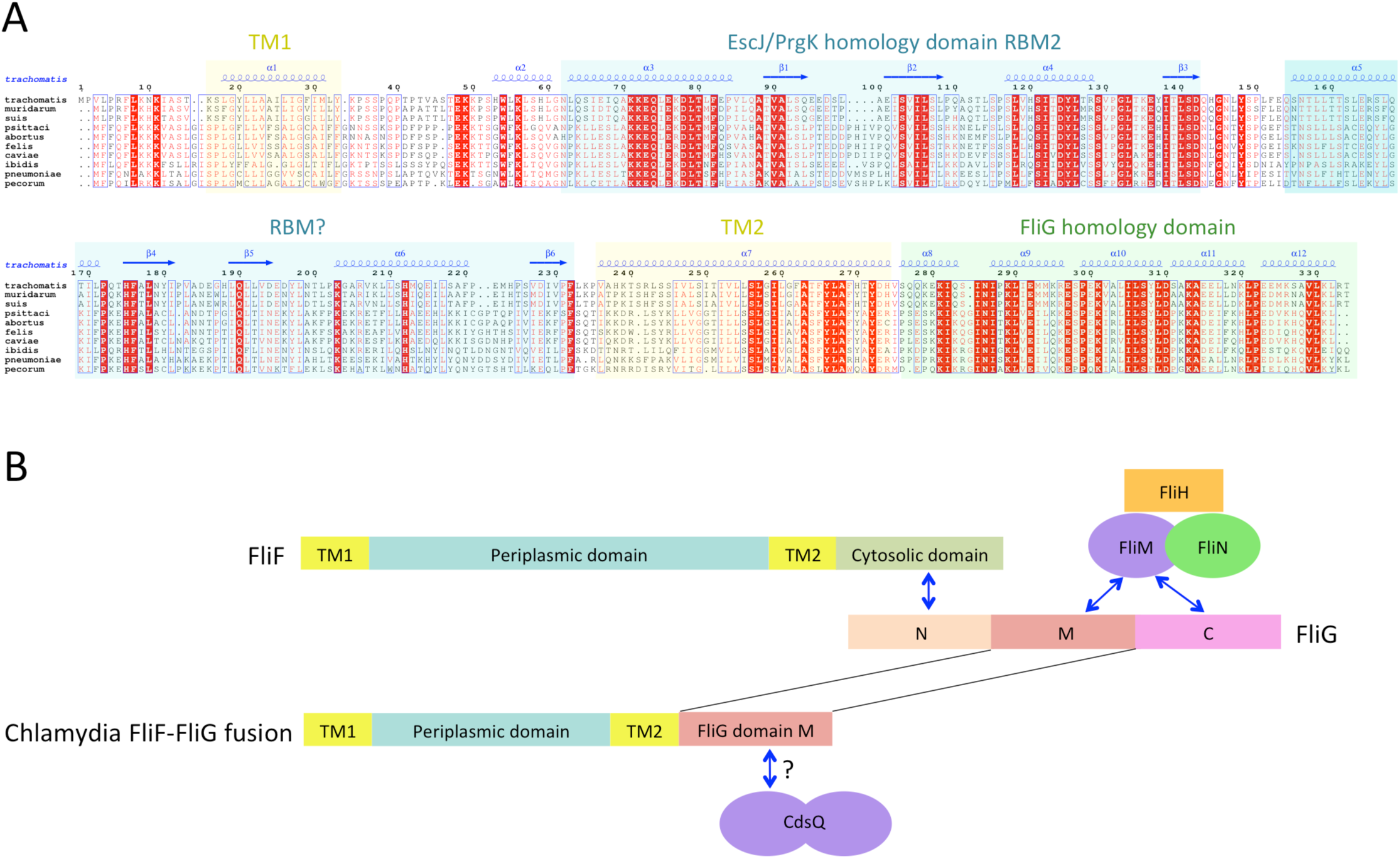
The *Chlamydia* FliF orthologue has unusual domain architecture. (A) Multiple sequence alignment of FliF sequences from the Chlamydiacae family (*C. trachomatis, C. muridarum, C. suis* from the genus *Chlamydia,* and *C. psittaci, C. abortus, C. felis, C. caviae, C. ibidis, C. pneumonia,* and *C. pecorum* from the genus *Chlamydophila.* Labeling is as in figure A, with the secondary structure prediction of the *C. trachomatis* orthologue shown at the top. (B) Schematic representation of FliF and its interaction with FliG (top), and of the *Chlamydia* FliF-FliG fusion (bottom).

In most orthologues, the C-terminus cytosolic region of FliF (residues 520–560) binds to the protein FliG^12,13^, a ~ 37 KDa protein possessing three domains, labeled N, M and C. FliF interacts with domain N, while both M and C bind to the stator (also known as C-ring) component FliM^25,26^, as illustrated on figure 3B. However, a sequence similarity search revealed that the in the *Chlamydia* FliF, the cytosolic region is actually homologous to the M region of FliG (not shown), revealing that in this species the protein is in fact a FliF-FliG fusion. *Chlamydia* is not thought be a flagellated bacterium, and possesses only a few flagellar genes, namely FliF, FliL and FlhA. It does however possess a functional T3SS that is essential for virulence^27^, and it has been shown that *Chlamidia* flagellar proteins interact with components of its T3SS^28^. FliM (and FliN, another C-ring component) is homologous to the *Chlamydia* T3SS component CdsQ (SpaO in *Salmonella*)^29^. It is therefore possible that the FliF-FliG fusion interacts with CdsQ (Figure 3B). This remains to be verified experimentally.

Interestingly, a FliF-FliG fusion is not entirely unprecedented, as two *S. typhimurium* strains containing such fusions have been reported^30,31^. In both cases, the fusion does not impair flagellum assembly and rotation, but induces a bias in the rotation direction. Since it is not yet clear if the *Chlamydia* FliF-FliG fusion protein is part of a proto-flagellum, or contributes to the T3SS, the implications for this observation is not clear.

### Structural modeling of the PrgK homology domain

Structures of the periplasmic domains of both EscJ^32,33^ and PrgK^19^ have been reported. Exploiting this information, a structural model for FliF was generated using the prototypical *Salmonella typhymurium* FliF sequence. Despite the predicted structural homology, the sequence conservation between FliF and PrgK is low (Figure 4A). I therefore employed a secondary structure alignment-based procedure for modeling the PrgK homology domain, which indicated that EscJ is the best modeling template, for the PrgK-homology region spanning residues 50 to 221. However, the relative orientation of the two RBMs in FliF is not known, and may differ from that of PrgK and EscJ. I therefore modeled the two RBMs independently, and refined the obtained models with Rosetta (see Materials and Methods for details). As shown on figure 4B and 4C, both RBMs converged to a local energy minimum within an RMSD of 1 Å to the lowest-energy model in the refinement procedure. The models possess good geometry (table 2), and their overall architecture (Figure 4D and 4E) is expectedly similar to that of EscJ and PrgK.

**Figure 4:**
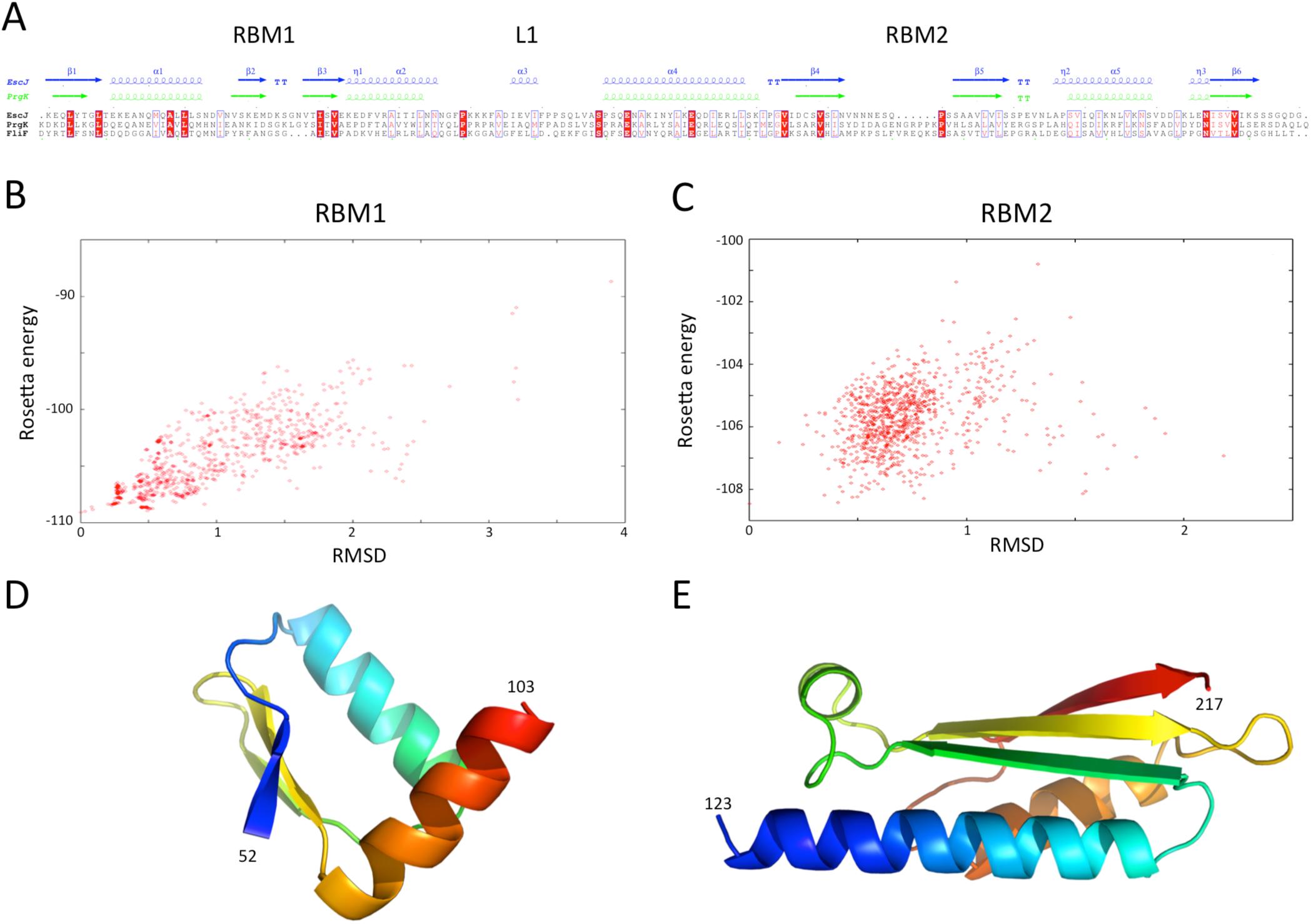
Modeling of the *S. typhimurium* FliF PrgK homology domain. (A) Sequence alignment of the EscJ and PrgK periplasmic domain sequences, with that of the PrgK homology domain of FliF (residues 50–221). Secondary structure elements for EscJ (PDB ID: 1YJ7) and PrgK (PDB ID: 3J6D) are shown at the top, in blue and green respectively. (B) and (C) Energy plot for the refinement of the FliF RBM1 and RBM2. The RMSD values are computed for all atoms, relative to the lowest-energy model. (D) and (E) Cartoon representation of the lowest-energy models for the FliF RBM1 and RBM2, with rainbow coloring indicating N- to C-termini.

**Table 2:** Geometry validation scores for the structural models of the three FliF RBMs.

|  | RBM1 | RBM2 | RBM3 |
| --- | --- | --- | --- |
| Ramachandran favored (%) | 98.0 | 97.8 | 98.9 |
| Ramachandran allowed (%) | 0.0 | 1.1 | 1.1 |
| Ramachandran disallowed (%) | 2.0 | 1.1 | 0.0 |
| Verify3D score | 0.46 | 0.38 | 0.31 |
| Procheck score | 0.39 | 0.26 | 0.20 |
| MolProbity Clashscore | 1.25 | 4.22 | 13.10 |
| Close contacts | 0 | 0 | 0 |
| RMSD bond angle (Å) | 1.5 | 2.6 | 1.8 |
| RMSD bond length (°) | 0.010 | 0.013 | 0.023 |

### The FliF-specific domain is a RBM

I next sought to generate a structural model for the FliF-specific domain (residues 228–443). As mentioned above, secondary structure prediction indicated that residues 228–309 and residues 356–443 possess defined structure, while residues 310–355 are predicted as intrinsically disordered (Fig 2). I therefore hypothesized that the FliF-specific region consists of two globular domains (D1 and D2) separated by a flexible linker. I attempted to identify structural homologues to these two regions. D1 shows structural homology to EscJ, although only for the first two structural elements (residues 229–268, Supplementary figure 2A). Similarly, D2 shows structural similarity to the sporulation complex protein SpoIIIAH, that also possesses a RBM fold and has been proposed to oligomerize into ring structures ^34,35^ (Supplementary figure 2B). However, the structural similarity was limited to the last three structural elements (residues 386–436). Based on these observations, I postulated that the FliF-specific domain is a “split” RBM that possesses a large insert in the loop between the second and third strand (Figure 5A). A structural similarity search for the FliF-specific domain with this insert removed (FliF_228-443_ Δ274–378) confirmed overall structural similarity to both EscJ and SpoIIIAH (Supplementary figure 2C).

**Figure 5:**
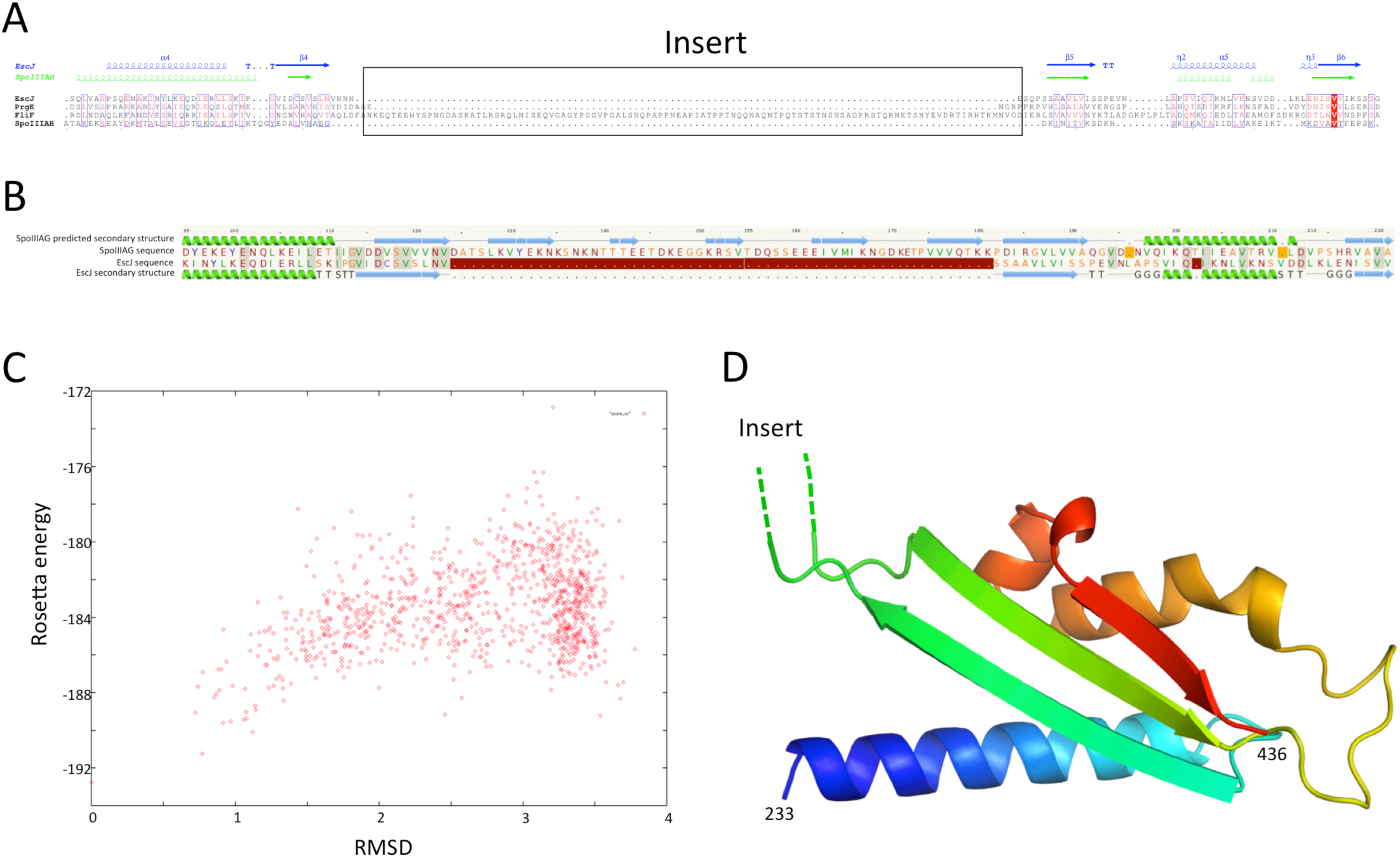
Modeling of the FliF-specific domain. (A) Sequence alignment of the EscJ RBM2, PrgK RBM2 and SPOIIIAH, with that of the FliF-specific domain (residues 228–439). Secondary structure elements for EscJ (PDB ID: 1YJ7) and SpoIIIAH (PDB ID: 3UZ0) are shown at the top, in blue and green respectively. The location of the FliF insert is indicated. (B) Secondary structure alignment between the *Bacillus subtilis* sporulation complex component SpoIIIAG and the T3SS component EscJ. Residues 124–181 of SpoIIIAG form an insert similar to that of the FliF RMB3. (C) Energy plot for the refinement of the FliF RBM3. The RMSD values are computed for all atoms, relative to the lowest-energy model. (D) Cartoon representation of the lowest-energy model for the FliF RBM3, with rainbow coloring indicating N- to C-termini. The location of the insert is indicated.

A “split” RBM is not unprecedented, as it is also predicted in the sporulation complex component SpoIIIAG (figure 5B). In both proteins the insert is predicted to possess four β-strands, which may indicate that it forms a small folded domain. Alternatively, it is possible that the insert adopts an extended loop conformation stabilized by β-strands.

In order to build an atomic model for the FliF RBM3, the EscJ-derived model for FliF_230–275_ was combined with the SpoIIIAH-derived model for FliF_380–440_, and refined with Rosetta. As shown on figure 5C, the RMSD to the lowest energy model is significantly higher than for the RBM1 and RBM2 models (around 3.5 Å for most decoys), which is perhaps expected, as the starting model was a presumably poorer, composite model. Nevertheless there is a clear energy funnel, indicative that the modeling procedure is converging. The obtained FliF RBM3 model, shown on figure 5D, possesses the canonical RBM fold, with good overall geometry (Table 2) and an elongated architecture as in the RBM2 of PrgK and in that of SpoIIIAH, with the insert between strands 1 and 2 located on one side of the structure.

### Localization of the FliF domains

Previous EM studies have shown that in isolation, FliF forms a doughnut-shaped oligomer with two side rings corresponding to the MS rings seen in the intact basal body^15,36^. The FliG-binding domain forms the cytoplasmic M ring, but little is known about the organization of the periplasmic domains. However, the organization of the T3SS basal body, and that of PrgK in particular, is well-characterized^18,19,37^. By comparing the EM reconstructions of the flagellar^15^ and T3SS^37^ basal body complexes, localization of the FliF RBMs was inferred. As shown on figure 6A, density attributed to the two globular domains of PrgK has clear equivalence in the flagellum, suggesting this location corresponds to the PrgK homology domain of FliF. EM map density for the flagellum S-ring corresponds to the T3SS protein PrgH localization, and is also present in purified FliF^36^, confirming that it is not attributed to another flagellar component. The periplasmic region of PrgH is composed of three domains with an RBM fold^18,20^, including the region corresponding to FliF density, suggesting that in the flagellum, this density can be attributed to the FliF-specific domain, which also possesses an RBM fold.

**Figure 6:**
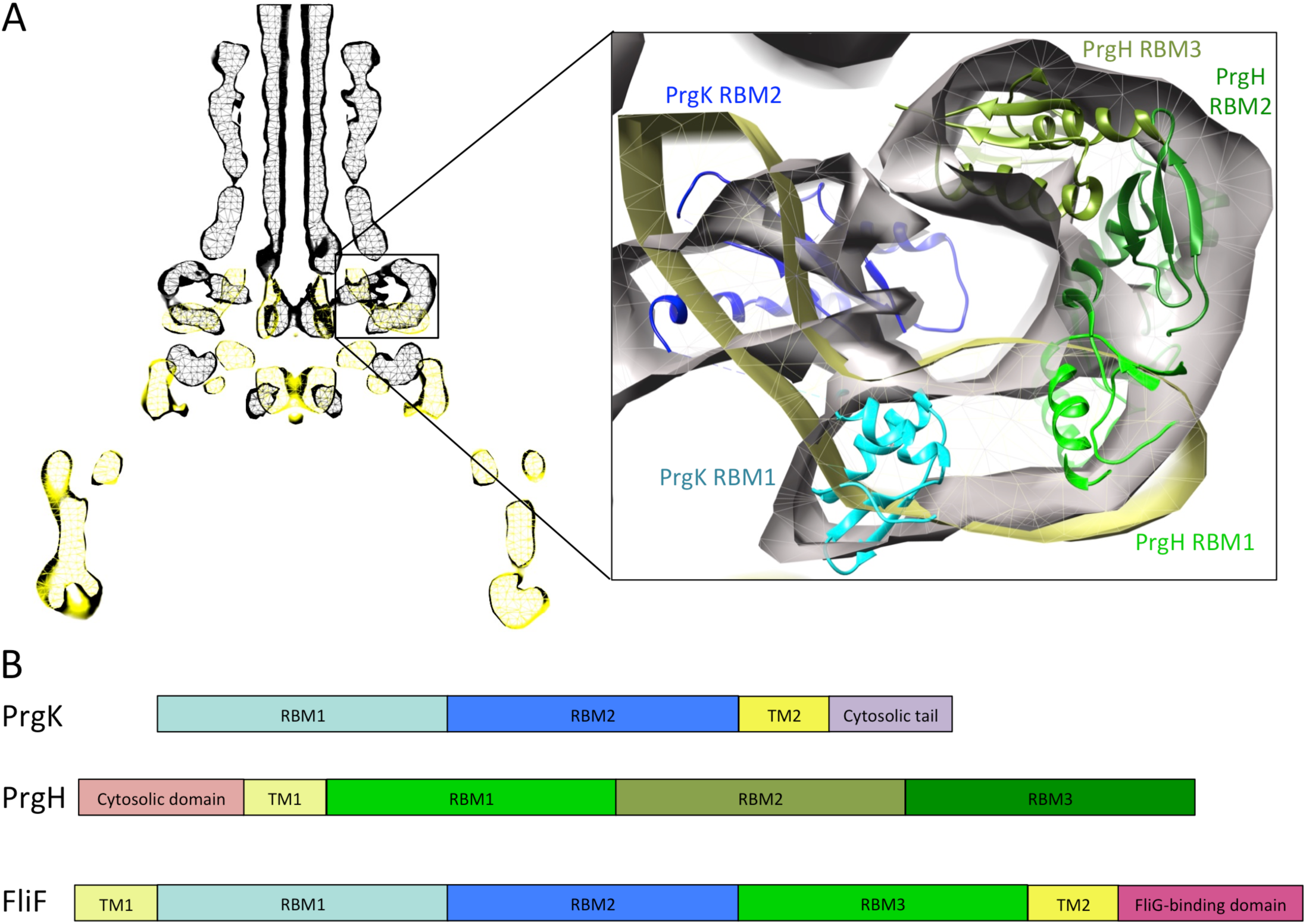
Comparison of the flagellar and T3SS basal body. (A) EM maps of the *S. typhimurium* T3SS basal body (EMBD ID: 1875) in black, overlaid on that of the *S. typhimurium* flagellum basal body (EMBD ID: 1887) in yellow. A close-up view of the S-ring region of the flagellum is shown on the left, with the PrgK/PrgH ring model (PDB ID 3J6D) docked in the T3SS model. The PrgK RBM1 is in cyan, PrgK RBM2 in blue, and the three PrgH RBMs are in green. (B) Schematic representation of PrgK, PrgH and FliF, with the TMs in yellow, and the RBMs colored as in (A) for PrgK and PrgH. Corresponding domains are indicated in FliF.

Based on these observations, I propose that FliF is akin to a PrgK-PrgH fusion (figure 6B), including the two RBMs of PrgK and its C-terminal TM helix, and the N-terminal TM and first RBM from PrgH. This likely explains a major difference between flagellar basal body, where FliF assembles on its own in the inner membrane, and T3SS basal body, where co-expression of PrgK and PrgH is required^38^.

### Modeling of the FliF oligomer

I next exploited the domain localization proposed above, the structural models of the three FliF RBMs, and the previously determined EM map^14^, to generate a model for the FliF oligomer. To that end, I positioned all three domains so that their termini point towards the correct region of the map, and for RBM1 and RBM2 in an orientation agreeing with the corresponding PrgK oligomers. Figure 7A shows that RBM1 is located near the inner-membrane region, while RBM2 forms the neck of the structure. Finally, the RBM3 is located in the region of density forming the S-ring. The insert in RBM3 points towards the lumen of the ring, and could correspond to the gate density observed in the FliF EM structure^14,36^.

**Figure 7:**
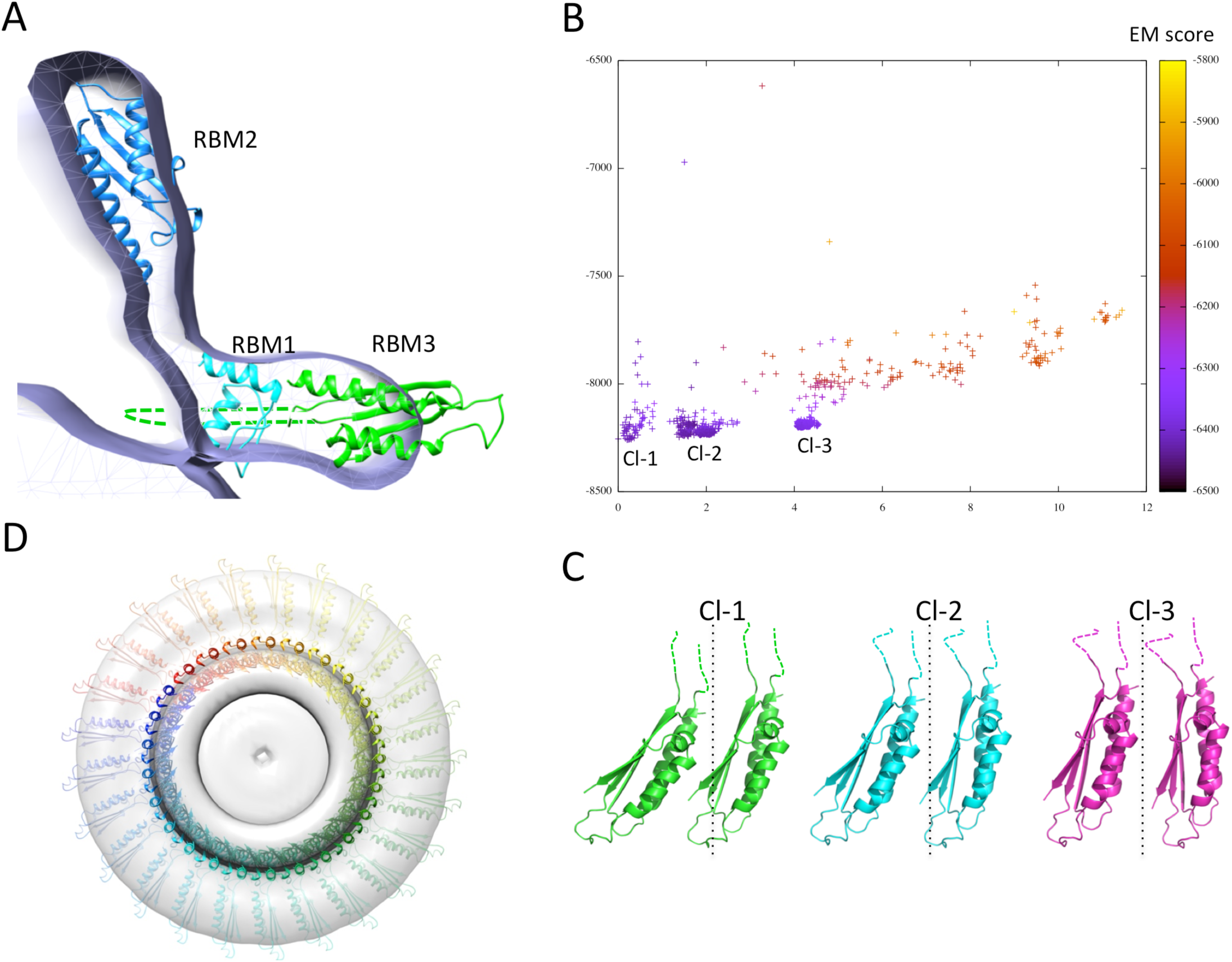
Modeling of the FliF oligomer. [A] Docking of the three FliF RBM modls in the FliF EM map. The domains are colored as in B. [B] Energy plot for the EM-guided symmetry docking procedure of the FliF RBM3. The RMSDs are computed for backbone atoms of the entire modeled 24mer complex, relative to the lowest-energy model, and color-coded depending on the fit to EM map. Three clusters of low-energy models were identified (Cl-1, Cl-2 and Cl-3], with two adjacent molecules for each cluster shown in (C). The 25-mer radius axis is represented by a dotted line. (D) 25-mer model of the FliF periplasmic region, viewed from the top.

I next attempted to generate a model for the FliF oligomer. I applied the previously described EM-guided modeling procedure^18^ to all three domains individually, using the FliF map as a restraint, and applying 24-, 25- or 26-fold symmetry. In most cases the procedure led to ring models that were too large for the EM map (not shown). This is unsurprising, as it had been reported previously that the procedure requires experimental structures (as opposed to homology models) to converge^18^. The one exception was the modeling of the RBM3, using 25-fold symmetry. This procedure generated three distinct clusters of models with clear low-energy funnels and good fit to EM density (Figure 7B). Close inspection of the clusters reveals that they correspond to similar oligomerization modes and use the same interface, but with a slightly different angle between the subunits (Figure 7C). I therefore exploited this 25-mer mode for the RBM3 to generate a complete 25-mer model for FliF. The corresponding model, shown in figure 7D and included in the supplementary material, matches well with the cryo-EM structure, with only a few loop regions of RBM2 and RBM3 located outside of the density. Further refinement of the model, including experimentally determined RBM1, RBM2 and RBM3 atomic structures, as well as a higher resolution EM map of FliF and of the intact flagellum basal body will be required to further refine this model.

## CONCLUSION

In this study I have demonstrated that the periplasmic region of most FliF orthologues consists of three globular domains possessing the canonical RBM motif. One exception is the Chlamydia FliF paralogue, which possesses only two of these, and has a FliG domain fusion at its C-terminus. By comparison with the T3SS basal body, I also propose the novel concept that FliF is akin to a fusion of PrgK and PrgH. Finally I have combined this information to propose a model for the oligomeric arrangement of the periplasmic region of FliF. Further experimental validation will be required to confirm these results, and to refine the FliF model.

These results shed new lights on the architecture and evolution of the flagellum MS ring. Specifically, the domain organization of FliF highlights similarities with the T3SS, but also with the bacterial sporulation complex. The proposed concept that FliF corresponds to a fusion of PrgK and PrgH likely explains why FliF can assemble spontaneously, while PrgK and PrgH require coexpression. In addition, the identification of a Chlamydia FliF-FliG fusion suggests that this may correspond to an ancestral complex.

## MATERIALS AND METHODS

### Sequence mining, analysis and alignment

All protein sequences were identified with the National Centre for Biotechnology Information protein database RefSeq^39^. Multiple sequence alignments were generated with ClustalW^40^ using default parameters. Alignment figures were produced with ESPript^41^.

Secondary structure elements, signal sequences, transmembrane helices and structural homologues were predicted with the PSIPRED server^42^. Protein sequence identity was calculated with the Needleman-Wunsch algorithm on the EBI server^43^, using default parameters.

### Modeling and refinement

Structure-based alignment and initial models were obtained with Phyre^44^, and the models were further refined by performing 1000 cycles of the Relax procedure^45^ in Rosetta 3.4^46^, using the following flags:

-database ~rosetta/rosetta_database
-in:file:s input.pdb
-in:file:fullatom
-relax:thorough
-nstruct 1000
-out:file:silent relax.silent

The geometry of the obtained models was analyzed with the PSVS suite^47^.

### EM map docking and symmetry modeling

The flagellum, T3SS, and FliF EM maps (EMDB ID 1887, 1875 and personal communication from K. Namba), were docked with the MatchMaker tool in Chimera^48^. Models of the FliF RBMs were placed in their putative location of the FliF EM map manually using Chimera.

For the EM-guided symmetry modeling of the RBM3, 24-fold, 25-fold and 26-fold symmetry definition files were generated, and used for the rigid-body step of the EM-guided symmetry modeling procedure described previously^18^. Briefly, the individual domains were manually placed in the corresponding region of the EM map, and 1000 rigid-body decoy models were generated with imposed symmetry and with a restraint for fit into the FliF EM map density, at 22 Å. The obtained models were isolated with the Cluster procedure in Rosetta, and scores calculated with the Score procedure using the lowest-energy model as a template for RMS calculations.

The flags used for the modeling procedure are listed in the Supplementary Methods section.

## ACKNOWLEDGEMENTS

I acknowledge Dr Keiichi Namba and Dr Hirofumi Suzuki for providing the FliF EM map. I thank Dr Natalie Zeytuni and Dr Morgan Bye for critical comments on the manuscript. This work was supported by a UBC Centre for Blood Research post-doctoral transition grant.

